# Impact of Fragmentation on the Metapopulation Structure of wild olive

**DOI:** 10.64898/2026.04.30.721863

**Authors:** Arayaselassie Abebe, Ramiro D. Crego, Markus P. Eichhorn

## Abstract

Habitat fragmentation disrupts metapopulation dynamics by altering environmental conditions and constraining demographic processes critical for persistence and recruitment. In the dry Afromontane forests of northern Ethiopia, we investigated how natural and anthropogenic drivers affect seedlings, saplings, and mature tree dynamics of *Olea europaea* subsp. *cuspidata* across 34 patches. We used dynamic occurrence models to quantify effects of patch area, altitude, browsing, and disturbance. Our results indicate that high disturbance reduces seedling occurrence probability lower disturbance sites has seedling in 30% of survey plots, high disturbance would bring this down to 10% (median = −1.322, 95% CI: −2.703 to −0.283). Disturbance makes seedling less likely to persist, while large patch size help seedling persists (median = −0.93, 9 5 % CrI −1.87 – −0.02). For mature individuals, disturbance was the only significant predictor of occurrence probability, suggesting greater resistance to environmental and spatial variability compared to earlier life stages. These findings emphasize that while mature trees display resilience, the successful regeneration of *Olea europaea* is constrained by disturbance, but current level of browsing is not a threat. Management strategies for conservation should prioritise reducing disturbance through community engagement and forest stewardship to enhance regeneration potential and ensure long-term population viability.

## Introduction

Populations are seldom continuously distributed across space and are often structured as metapopulations (Bergerot et al., 2010; Opdam, 1991; Thompson et al., 2017). These spatial arrangements are exhibited more prominently in areas with fragmented landscapes (Aguilar et al., 2006; Forman., 1995; Kramer et al., 2008; Young et al., 1996). The expansion of agriculture with the consequential land clearing has resulted in the fragmentation of many forest ecosystems worldwide (Li et al., 2022). In such fragmented forests, the presence or absence of plant species within each forest patch is influenced by local factors such as soil conditions, microclimate, and abiotic interactions (Elias, 2016). The persistence of species within these landscapes depends on habitat quality, connectivity, and the dispersal ability of the species (Hanski, 1994). To maintain viable metapopulations, trees in fragmented landscapes require interconnected patches rather than isolated large areas due to complex source-to-sink dynamics influenced by dispersal, seed banks, habitat management, pollinators, and climatic stressors (De Kort et al., 2018; Kramer et al., 2008). Habitat degradation intensifies edge effects, reduces genetic diversity, and disturbs species interactions, which can further increase connectivity thresholds for population persistence (Forman., 1995; Riva & Fahrig, 2023). Some populations may benefit from fragmentation alone (Haddad et al., 2015), but to improve connectivity, conservation strategies must give priority to corridors and stepping-stones. Where fragmented forest patches are threatened by deforestation and land degradation, even though remnant biodiversity offers the possibility of restoration, maintenance of connectivity is particularly critical (Abiyu et al., 2016; Cardelús et al., 2019; Teketay, 2001).

In northern Ethiopia, most forest patches remain around churches or on steep terrain which serve as refugia for species (Abebe et al., 2023). The Ethiopian Orthodox Tewahido Church (EOTC) forests have been recognised as a biodiversity hub by previous work in this region (Klepeis et al., 2016). Most of the church forest patches are grouped under the vegetation type of Afromontane forests (Kindu et al., 2022). These patches are often perceived not merely as ecological remnants, but as holy sanctuaries and refugia for both biodiversity and spiritual values. Their continued protection is deeply rooted in indigenous Christian theology, with rituals, prohibitions, and customary laws ensuring their survival (Bongers et al., 2006; Cardelús et al., 2019; Wassie et al., 2005). The land preservation by churches has resulted in a patchy distribution of remnant forests within a matrix of different land uses, predominantly agricultural and pastureland (Nesibu *et al*., 2022). The patches owned by the church can be relatively new, and not all of them are remnants of original vegetation. This illustrates how the landscape is ever-changing due to ecological processes and shifts in land use. From a socio-ecological standpoint, the system is especially compelling because of the coexistence of older and newer patches, providing insights into the interactions between ecological dynamics and human activity over time.

Patch formation influences the distribution and abundance of plant species with important cultural and economic values, including several narrow-range and endemic species. The wild olive (*Olea europaea* subsp. *cuspidata* [Wall. ex DC.] Ciferri), one of the six subspecies within the *Olea europaea* complex (Guerrero *et al*., 2016) is one such example. The species is a key indicator of Dry Afromontane Forest and is under significant threat due to over-harvesting (Aynekulu et al., 2009). The sub-species is endemic to and widely distributed across southern to north-eastern Africa, Southwest Asia and the arid regions of China’s Yunnan and Sichuan. *Olea europaea* subsp. *cuspidata* is a prominent late-successional species in the region (Aynekulu et al., 2009). *Olea europaea* is a highly valued tree in Ethiopia and worldwide for medicines (root, leaf, bark, and wood), construction, farm tools (handles and pins), firewood and cultural practices, which include fumigating food utensils and traditional drink preparation (Ourge et al., 2018). The mature trees of this species are presently confined to churches, protected areas and inaccessible areas of Northern Ethiopia (Abiyu et al., 2016).

In this study we investigated the metapopulation dynamics of *Olea* across a matrix of forest patches. Specifically, we examined: i) how do occurrence, persistence and recruitment of *Olea europaea* subsp. *cuspidata* seedlings vary across different seasons and forest patches, ii) what are the environmental and anthropogenic factors driving these patterns; and iii) what are the environmental and anthropogenic drivers of *Olea europaea* subsp. *cuspidata* occurrence for saplings and mature trees. Understanding the factors driving the establishment of new seedlings and the growth of *Olea* trees among patches is critical for ensuring the effective conservation of a crucial structural and cultural component of forest patches in northern Ethiopia.

## Methodology

### Study area

The study was conducted in the Northern Wollo Kobo district of Ethiopia, located between 12.194N-39.766E and 12.105N-39.784E (Fig. 1). This area covers approximately 12172 – 12180 km^2^, highland > 2400 m.a.s.l, mid-altitudes (1,250 – 2,400 m.a.s.l.), and lowlands (> 1250 m.a.s.l. The vegetation is primarily Dry Evergreen Afromontane Forest with Acacia (Vachellia spp.) woodlands found in the lowland areas. Annual rainfall is 2000 mm and is concentrated over a wet season lasting three months (July to September), followed by an extended dry season lasting nine months (October to June).

**Fig 1.**
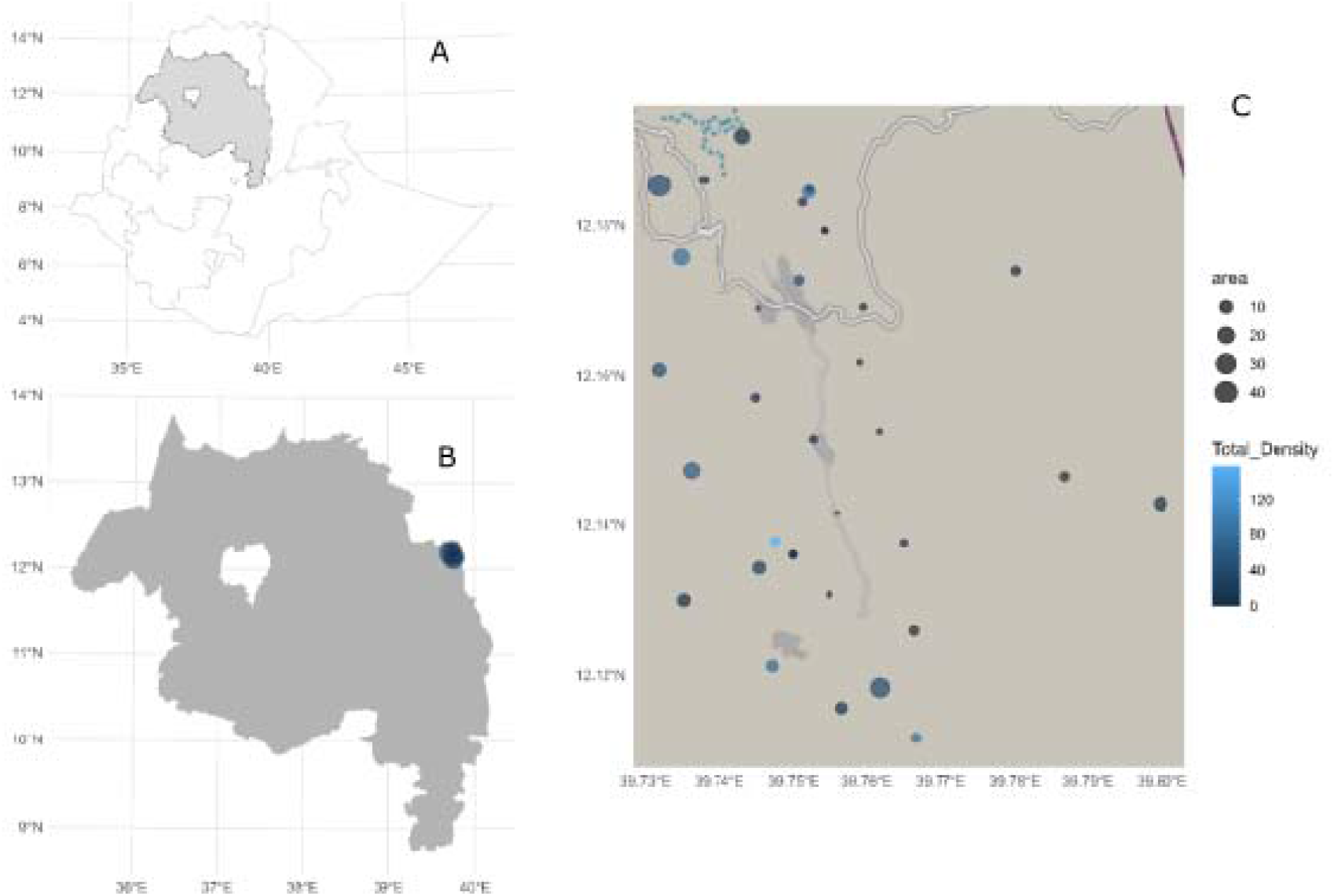
Study area map A. Ethiopia with regional administration. B Amhara regional state and C. Map showing the location of the forest patches surveyed in the Northern Ethiopia study area, illustrating each patch’s area and density *of Olea europaea* subsp. *cuspidata* seedling.

### Study design and data collection

We delineated habitat patches based on vegetation cover and patch size, adhering to the FAO forest classification guidelines for Ethiopia (FRA, 2015). The criteria states that any land spanning more than 0.5 ha with trees higher than 5 m and a canopy cover of more than 10 percent can be called a forest patch. We manually identified 34 patches from high resolution imagery available on Google Earth (Gorelick et al., 2017). For each patch, we first delimited the area by drawing polygons over Google Earth images. We then refined those polygons with waypoints collected on the ground along the edges of patches using a hand-held GPS unit (Garmin GPSMAP 65).

To understand the metapopulation dynamics of *Olea* across the annual cycle, we conducted vegetation sampling during February to March 2020 (first dry season), September to October 2020 (first wet season), March to April 2021 (second dry season), and September to October 2021 (second wet season). Transects of up to 400 m length were created across the interior of each forest patch. The number, location and length of transects varied depending on the topography, shape and size of each forest patch. The topography of the patch is related to the areas in which monasteries /churches have been set, either flat areas or mountain peaks. For patches located on flat ground, we placed the first transect in the eastern side of the patch, from which subsequent transects were placed in parallel, separated by 400 m. For patches located on peaks, we placed all transect starting points approximately 10 m from the establishment point (where the monastery is located), with each transect radiating towards the periphery such that they were separated by a minimum of 400 m at the patch edge. The size of the patch determined the total number of transects that were contained within each patch, resulting in a total of 116 transects (mean 3.11 and range 2--4). Within each 400-m transect we placed 20 × 20 m plots separated by 50 m. Within each plot, we placed a 5 × 5 m subplot at the centre and five 1 × 1 m subplots, one at the centre and four at the corners of the larger plot (Figure 2). This resulted in a mean number of plots per patch of 5.1 and range 3-15.

**Fig. 2.**
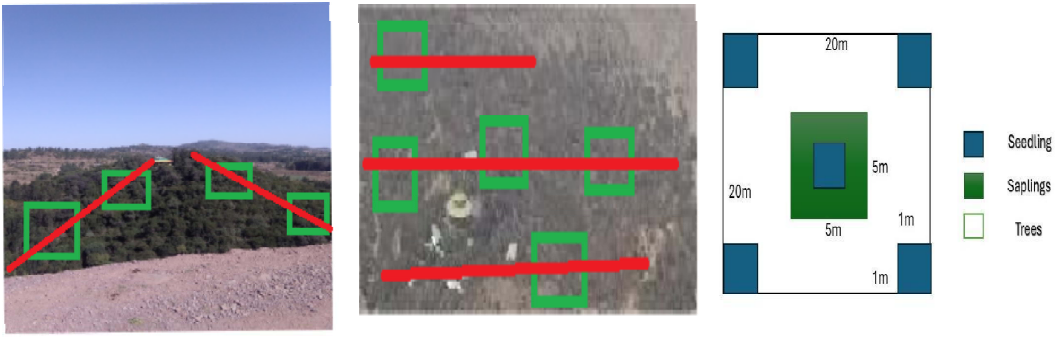
Schematic plots and transects layout for (A) transects used in hilltop areas, (B) transects in flat landscapes and (C) plot size and nested plot arrangement used to collect data for seedlings, saplings and mature trees of *Olea europaea* subsp. c*uspidata* in the study area. Plot representations on the figure are not to scale and are intended to illustrate the sampling design. (Photo credit, Arayaselassie Abebe 2022)

Within each plot we counted the total number of *Olea* seedlings (height <1.3 m) within each 1 × 1 m subplot, saplings (plant height >1.3 m and DBH <5 cm) within the 5 × 5 m subplot, and trees (plant height <1.3 m and DBH >5 cm) within the full 20 × 20 m plot. We measured the DBH of mature trees of *Olea* using a measuring tape. We collected a pressed sample of trees from each patch and confirmed the species at Addis Ababa University National Herbarium using the Flora of Ethiopia and Eritrea volume 1-8 (Hedberg et al., 2009). Plots and subplots were marked using white paint and stones for consistency of sampling during revisits across different seasons.

### Covariables

At each patch, we recorded covariables that could influence *Olea* abundance, occurrence, and recruitment. Area (ha) was recorded from legal documents. Altitude (m asl) is the mean plot elevation per patch. We measured disturbance on an ordinal scale. Low disturbance (1) refers to conditions where light thinning or minor natural events occur, maintaining a largely intact canopy cover of over 70%. Moderate disturbance (2) is characterised by partial clearance that reduces the canopy cover to 30% and 70%. High disturbance (3) involves clear-cutting, leading to canopy cover dropping below 30% (Turner,2010). Recurrent disturbances of patches include the establishment of roads, meeting places, clearings, illegally harvested trees, and clearing for graveyards around churches. These and other anthropogenic disturbances were included in the disturbance rate (Muluneh et al., 2021).

Browsing intensity was measured based on local abundance of livestock. We made a single assessment of browser abundance because livestock populations vary little over the timescales captured by the study (Cardelús et al., 2019;).We collected information about the associated community, including households and their livestock surrounding each patch. We acquired this data from two primary sources: the local administration and the Ethiopian Statistics Agency’s national census reports (SAE, 2022). When two churches or monasteries are close to each other, to avoid double counting the households and associated livestock, locality-based data was used with support from agricultural experts in the administration to determine the patch which they belong to and use. Kebeles are small administrative bodies composed of a collection of localities or rural villages with variable numbers of households influenced by migration and landscape. The believers are registered as members of each church, or “Atebya.” Households are subsequently categorized into local groupings termed “gote,” which refers to small villages and the church designated for their service. Households only hold customary rights for use of a single patch of forest and graze livestock within a defined common, which makes it possible to definitively assign households and livestock to each forest patch.

Finally, we included the age of each patch, which we determined using documents from local administrative bodies (Kittler, 2016). The age of each church forest patch is based on the establishment year of the church. Churches are built in the middle of forest patches and therefore the age also indicates the time of isolation of the patch. In the case of protected and restored areas, we used the legal date of closure as a starting point (Klepeis et al., 2016).

### Data analysis

#### Metapopulation modelling framework

To understand the seasonal dynamics of *Olea europea* ssp. *cuspidata* seedlings across the fragmented landscape we applied a modified dynamic occurrence model (an occurrence model for which we assumed perfect detection) that incorporated occurrence probability for season one, and for which subsequent occurrence probability for the other seasons was estimated by incorporating persistence (opposite to extinction) and recruitment probabilities from season to season. This type of dynamic model is particularly well-suited for capturing persistence and recruitment processes at each patch in fragmented systems (MacKenzie et al., 2003).

In the specification of the model, we modelled occurrence probability at each patch *i* in season one (ψ_i,1_), the first dry season, as an outcome of a Bernoulli trial:

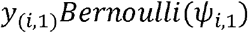

and for the following seasons, occurrence probability at each patch *i* at a season *t* is also the outcome of a Bernoulli trial of the sum of the product of probability of persistence (φ; probability that an occupied patch remains occupied) and the occurrence status at *t-1*, plus the product of the probability of recruitment (γ; probability that an unoccupied site becomes occupied) and the probability that the site was not occupied at time *t-1*.

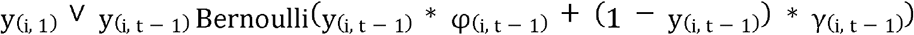

We allowed persistence and recruitment probabilities to vary across seasons given that we would expect these parameters to differ in transitions from wet to dry or dry to wet seasons.

We then modelled probability of occurrence at season one, probability of persistence and probability of recruitment as a function of patch area, disturbance index, altitude and number of browsers using a logit link function.

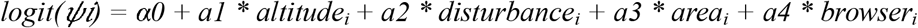

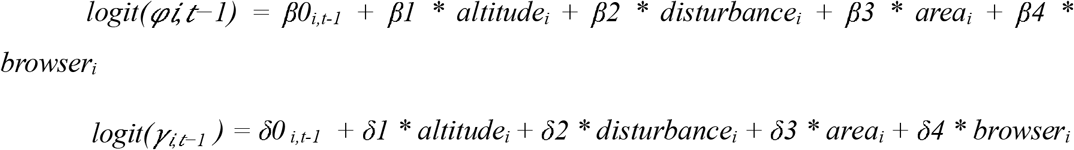

To understand occurrence probability of saplings and mature trees, we fit a logistic regression using a logit link function to incorporate patch area, disturbance index, altitude and number of browsers as covariables. Variance inflation factors (VIF) among covariables were < **1.8, ensuring no collinearity problems**.

We fit all models in a Bayesian framework through the NIMBLE package v1.3.0 (de Valpine et al., 2017, 2022) in R programming language v4.4.1 (R Core Team 2024). We set uniform priors on the logit scale for intercepts and weakly informative Normal distributions (μ = 0, σ = 1.5) for all other parameter coefficients. We ran three Markov Chain Monte Carlo (MCMC) cycles for 60,000 iterations, discarding the first 20,000 and thinning by 40, resulting in 3,000 total samples. Convergence was checked by visually inspecting the trace plots of each parameter and ensuring an effective sample size > 1,000 and a Gelman-Rubin statistic < 1.01.

#### Tree Growth Dynamics and Patch Age Effects

To assess long-term ecological dynamics, we modelled the relationship between patch age and diameter at breast height (DBH) of mature *Olea* individuals. We applied a simple linear regression using standardised patch age as the independent variable and mean DBH per patch as the dependent variable.

## Results

Of the 34 forest patches surveyed over four seasons, 23 (67.6%) contained at least one *Olea europaea* subsp. *cuspidata* individual in at least one sampling season.

### Persistence, Recruitment, and Occurrence

Posterior estimates indicated that disturbance had a strong negative effect on seedling occurrence. Seedling occurrence probability decreased with increasing disturbance (median = –1.322, 95% CrI: –2.703 to –0.283) with occurrence probability at lower disturbance of 39%whereas at higher disturbance it fell to 4.5%. There was also a decrease in occurrence with increasing patch area (median = –0.273, 95% CrI: –1.358 to 1.070), but credible intervals overlapped zero, whereas seedling occurrence probability was not affected by altitude (median = 0.019, 95% CrI: –0.975 to 1.102) (Fig. 3). Number of browsers also showed a weak negative effect on occurrence but with CrI overlapping zero

**Fig 3.**
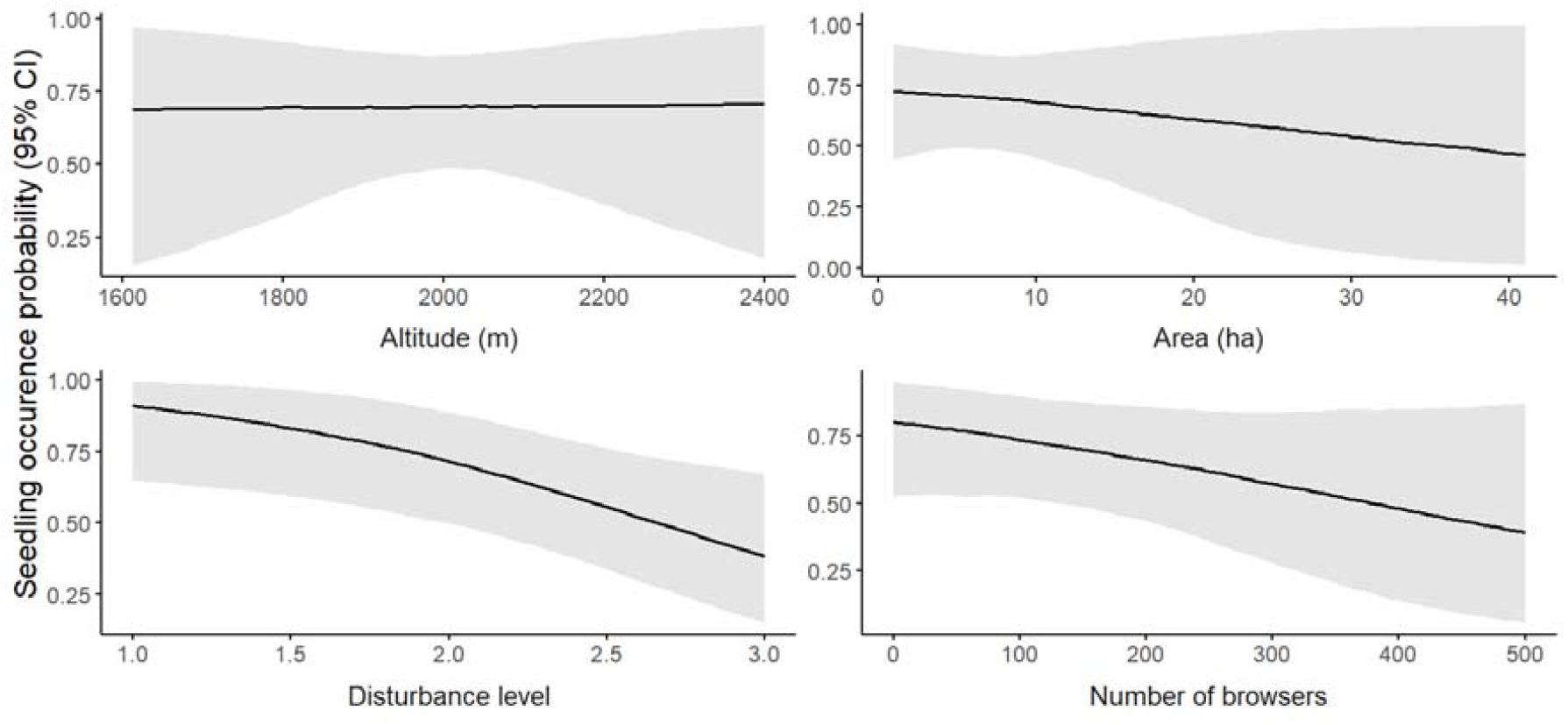
Median occurrence probability and 95% credible intervals (CrI) as a function of altitude, patch area, disturbance, and browser abundance of *Olea europaea* subsp. *cuspidata*. seedlings across 34 forest patches in northern Wollo.

In the first transition from wet to dry season, seedling persistence probability increased with increasing browser abundance (median = 0.90, 95 % CrI 0.54 – 0.99) and with increasing disturbance levels (median = 0.70, 95% CrI 0.34 – 0.92), while altitude showed a weaker but still positive effect (median =0.60, 95% CrI 0.29 – 0.86) (Fig.4).

**Fig 4.**
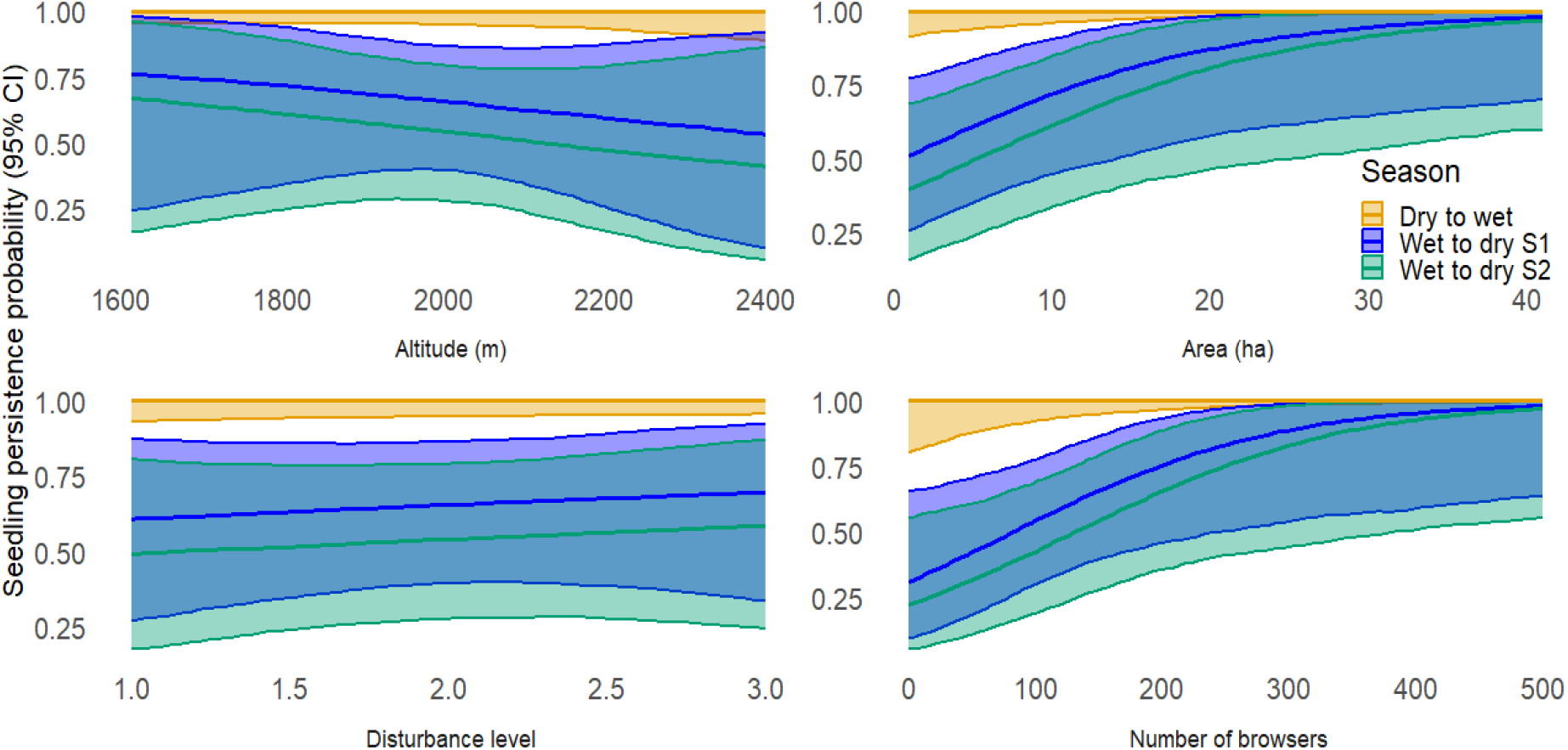
Seedling persistence probability as a function of altitude, patch area, disturbance, and browser abundance across three seasonal transitions. Error bars indicate 95% credible intervals.

Larger patches consistently increased persistence probability (median = 0.88, 95 % CrI 0.11 – 1.72). During the dry to wet season transition, persistence probability reached a maximum, with all three environmental drivers approaching unity: altitude (medium = 0.99, 95% CrI 0.943 – 1.00), browsing (median =1.00, 95% CrI 0.984 – 1.00), and disturbance (median =1.00, 95% CrI 0.958 – 1.00).

The second wet to dry season shift had similar responses to the first. Browsers (median = 0.84, 95% CrI 0.45 – 0.98) and disturbance (median =0.59, 95% CrI 0.24 – 0.87) still raised persistence probability, whereas the effect of altitude decreased but remained positive (median =0.49, 95% CrI 0.19 – 0.78). Patch area has a consistently positive effect on persistence probability, with larger patches tending to maintain seedling presence from one season to the next, and this relationship did not vary between wet to dry or dry to wet transitions (median = 0.88, 95% CrI 0.11 – 1.72).

Recruitment from the first wet to dry season transition was very low (median = 0.01, 95 % CrI < 0.001 – 0.15 %) indicating that almost no new seedlings became established. The model identified negative impacts of both disturbance (median = –0.69, 95 % CrI –1.84 – 0.38) and browser abundance (median = –0.62, 95% CrI –1.82 – 0.37), but credible intervals spanned zero. Effects of patch area (median = 0.36, 95% CrI–1.32 – 2.24) and altitude (median = 0.35, 95% CrI –0.92 – 1.79) on recruitment were not statistically significant, as their 95% credible intervals overlapped zero, providing no clear evidence that either variable influenced recruitment.

Sites with more disturbance and heavier browsing has lower recruitment, while larger patch area and higher altitude showed only slight increases in recruitment. During the second wet to dry season transition, recruitment was virtually zero. This pattern showed the first wet to dry shift; disturbance and browsing were negative, area and altitude were at best weakly positive, but the overall baseline was so low that recruitment was effectively absent in that season (Fig.5).

The second wet to dry transition again showed almost zero recruitment (0.01%). The covariate effects parallelled those of the first wet to dry shift with negative effects of disturbance and browsing, equivocal effects of area and altitude, but overall, an extremely low intercept, leaving recruitment effectively absent during this season (Fig. 5).

**Fig. 5.**
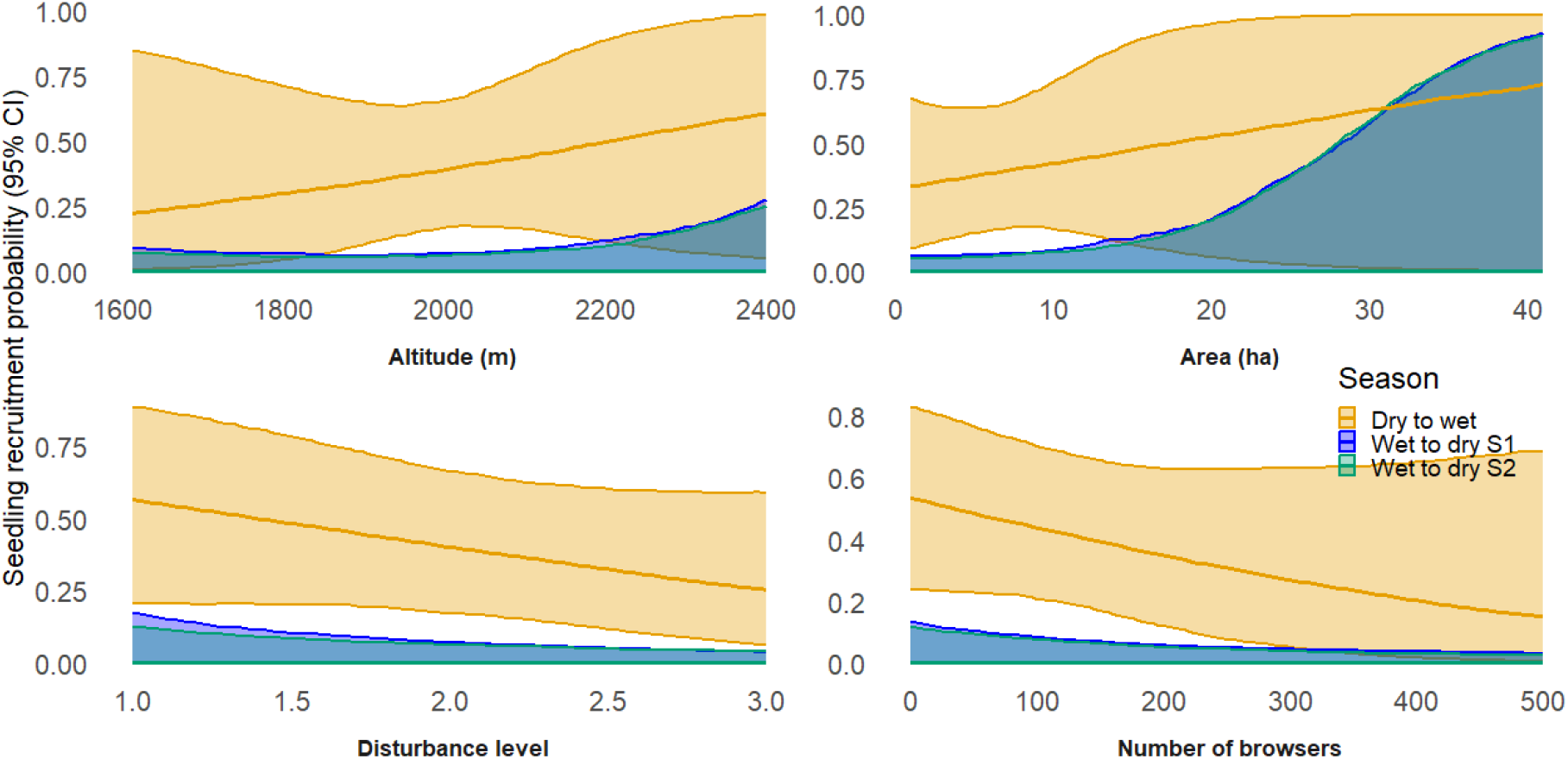
Recruitment probability in relation to altitude, patch area, disturbance and browser abundance across three seasonal transitions in forest patches of northern Ethiopia.

### Effect of environmental and anthropogenic effects on abundance of *Olea* in different growth stages

Sapling occurrence increased with patch area (β = 2.61; 95% CI: 0.61 to 4.99) and decreased with disturbance (β = −3.54; 95% CI: −5.00 to −1.86). Altitude and browsing effects overlapped zero (Fig. 6). For mature individuals, disturbance was the primary factor negatively affecting the abundance of mature *Olea europaea* subsp. *cuspidata* trees. The significant negative relationship (β = −1.51; 95% CI: −2.80 to −0.35) indicates that as disturbance intensity increases the likelihood of matured tree occurrence decreases. In contrast, altitude, patch area and browsers were not significant (Fig. 7).

**Fig 6.**
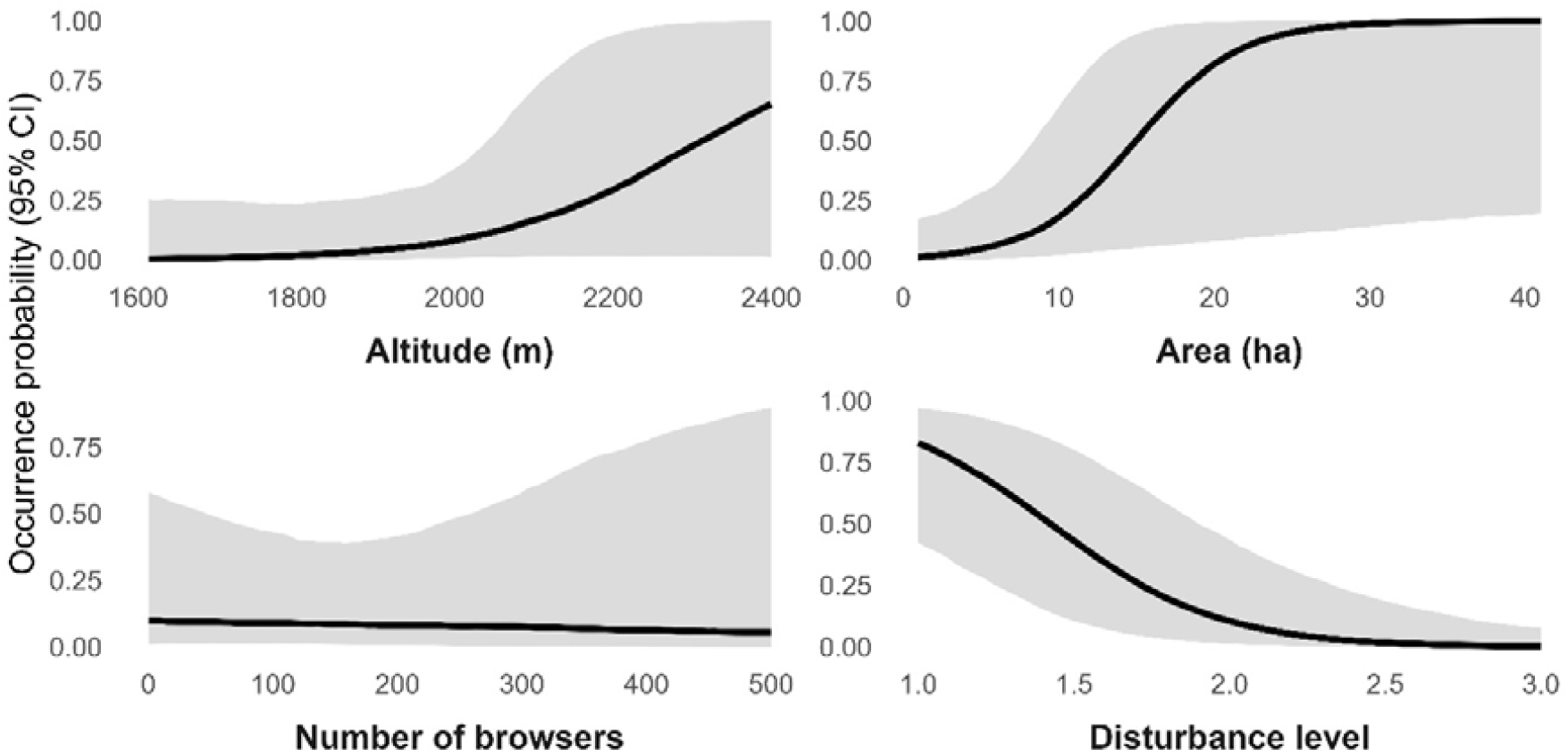
Effect of altitude, patch area, browsers, and disturbance on *Olea europaea* subsp. *cuspidata* saplings across forest patches of northern Wollo Ethiopia.

**Fig.7.**
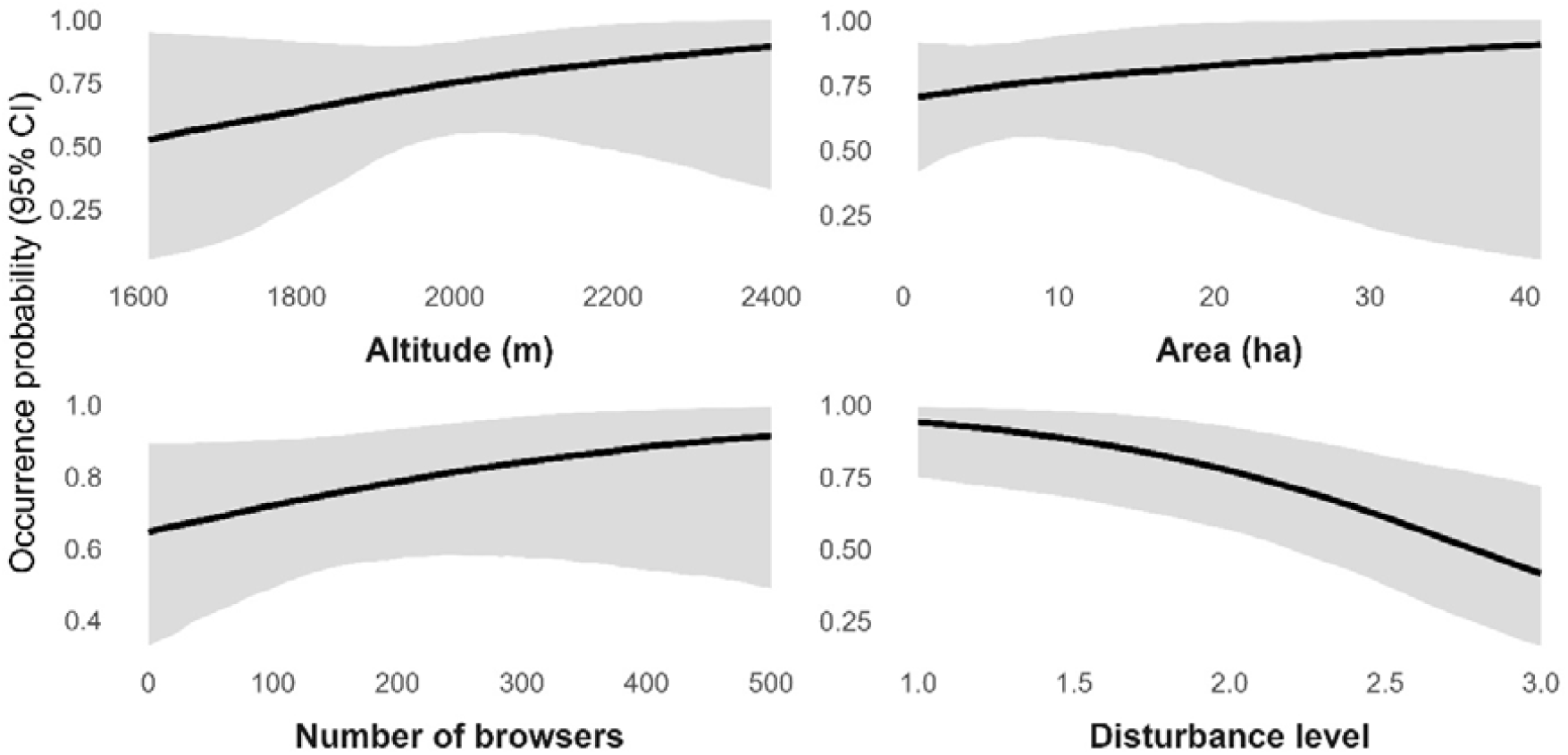
Effects of altitude, patch area, browser abundance, and disturbance on *Olea europaea* subsp. *cuspidata* mature trees across forest patches of northern Wollo Ethiopia.

### Mature tree structure proxies

Patch age was strongly and positively related to the diameter at breast height (DBH) of mature trees (R^2^ = 0.886, p < 0.001), indicating that disturbance reduces the size of matured tree, whereas stand age promotes structural development.

## Discussion

Within a fragmented forest habitat in northern Ethiopia, two thirds of the 34 patches contained *Olea europea* ssp. *cuspidata* at least once over the four seasonal transitions. Occurrence patterns were consistent with metapopulation processes in a fragmented, mixed biocultural landscape. Disturbance reduces seedling and sapling occurrence, while saplings benefit from large patch areas, implying demographic recovery in large fragments. Mature tree presence is mainly constrained by disturbance, suggesting the legacy impacts of human pressure despite adult resilience.

The seasonal patterns show that higher disturbance lowers initial occurrence by 66% while heavy browsing decreases recruitment odds by nearly half (β = −0.66, −1.82 – 0.37). The drop in seedling recruitment under high disturbance and browsing intensity is consistent with trends seen in Amhara and Tigray regions, where extensive ungulate browsing decreased regeneration success by as much as 75% in open-access forests (Teketay, 2005; Temesgen & Warkineh, 2020). According to Angassa & Oba (2010), open access forests experience 3--5 times higher livestock density than church forests, directly correlating with 60-80% lower seedling survival rates. Canopy dependent species like *Juniperus procera* and *Olea europaea* show >90% recruitment failure under combined browsing and trampling (Aerts et al., 2007, 2016).

Seedling establishment occurs at the end of the rainy season in July/August, as indicated by a recruitment increase. The results also indicate that wet seasons have a higher probability of recruitment (β□ = −0.42) with change in presence of *Olea* seedling from 32.4% to 58.8%. Seedling persistence is near constant at 0.99 (0.943 - 1.00) before decreasing to 0.49-0.60 in the following dry season. Seasonal variation in soil moisture (θv) and the rainfall threshold at which seedling wilting occurs influence seedling persistence (Pinto et al., 2016). Field observations and controlled exclosures experiments in urban forest in Addis Ababa have shown that the transition from dry to wet conditions shows 90% seedling persistence, underscoring the critical role of water availability in seedling survival (Tenkir et al., 2024). Conversely, prolonged dry spells or the transition from wet to dry conditions often lead to a 15–25% decline in seedling biomass and survival, particularly among drought-sensitive species (Bekele, 2005; Wassie et al 2009). Such seasonal dynamics suggest that early establishment during favourable moisture conditions is essential to buffer against later water stress (Teketay, 2005).

Larger patches increased persistence, but that advantage disappeared once browser density exceeded 15 km^-2^, at which point anticipated recruitment fell below 0.05 independent of patch size. Intense anthropogenic disturbance such as logging and fuel wood collection can reduce *Olea europaea* seedling survival by a factor of six, as documented in northern Ethiopia’s Hugumburda forest (Aerts et al., 2007). In contrast, a decade-long herbivore manipulation at the Kenya long term exclosures experiment showed that moderate grazing suppresses grasses and fire intensity, boosting woody recruit persistence by roughly 30% (Wegasie et al., 2021). In addition, high browsing pressure has been limiting the transition of seedlings to saplings. In fragments where ungulate density exceeds c.15 km^-2^, browsers remove upon to 70% of current year shoots, creating a demographic bottleneck that overrides the limited buffering capacity of the larger habitat cores (; Wassie et al., 2009). Establishing low-browser thickets and corridor strips has been shown to double recruitment rate while simultaneously diluting local herbivore impacts by dissipating browser pressure across a broader network (De Kort et al., 2018; Riva & Fahrig, 2023).

Patch age and DBH have a positive, nearly deterministic relationship (R^2^ = 0.886), which highlights how crucial time-protected legacy forests are to preserving mature structural traits. This is in line with (Nyssen et al., 2014), who pointed out that “demographic shadows” are produced by past land use and sporadic disturbances, upsetting the age structure and succession of forests. These patterns support earlier research that suggest protection and active structural rehabilitation are necessary for restoration, especially in areas recovering from past overexploitation (Aynekulu et al., 2009; Wassie et al., 2009).

Our results are consistent with observations made in tropical dry forests around the world, such as the Caatinga in Brazil, the Miombo woodlands of East Africa, and the teak forests of Southeast Asia, where anthropogenic stress and fragmentation most frequently affect the early stages of regeneration (Bongers et al., 2006; Laurance et al., 2018; Newmark, 2008). Because of their proximity to developing agricultural frontiers and ongoing underrepresentation in conservation planning, Afromontane ecosystems are under considerable stress. Only <5% of Afromontane church forests are formally protected versus 14.7% of Ethiopia as a whole (Aerts et al., 2016).

The Ethiopian Orthodox Tewahido Church (EOTC) forests’ status as holy natural sites is one of the characteristics that distinguish them from the broader Ethiopian forest matrix. These patches are safeguarded by centuries-old religious practices, convictions, and canonical stewardship rather than by official legal means (Klepeis et al., 2016; Wassie et al., 2005). With higher mature tree survival and more reliable grazing protection, our results validate that spiritually governed forest patches frequently function as de facto conservation zones.

In Ethiopia and elsewhere, the idea of spiritual ecosystem services, the role of forests in ritual, identity, and continuity should be recognized as a fundamental component of conservation. According to Cardelús et al. (2019), sacred forests serve as resilience nodes in socio-ecological systems, providing examples of biocultural adaptation in which ecology and theology support conservation. Co-management frameworks based on biodiversity, tradition, and belief can be promoted by combining ecological science and theological stewardship. Supporting faith-based restoration theology, church-led exclusion zones, and the transmission of sacred knowledge across generations, for example, may sustain conservation outcomes even in areas with limited resources. Such approaches are essential to the long-term persistence of culturally important tree species like *Olea europaea* at a landscape scale through creating the necessary conditions for a sustainable metapopulation.

Our results point out the potential management levers: (i) reduce anthropogenic disturbance within patch cores (limit clearing, fuelwood removal, and graveyard expansion), (ii) manage browsing/grazing via seasonal exclusion specially during dry to wet season or rotation schemes, and (iii) protect and enlarge patches as regeneration anchors. In church forest contexts, stewardship and community norms can operationalise these measures with comparatively low cost. This study spans four seasonal surveys across 34 patches which, while sufficient for estimating intra annual transitions, is not able to resolve long term metapopulation dynamics in slow-growing trees. We therefore report seasonal transition parameters and stationary cycle illustrations rater than forecasts which can only be determined through long-term monitoring.

## Conclusion

Disturbance and grazing undermine the persistence and recruitment of *Olea europaea* subsp. *cuspidata* across fragmented forest landscapes in northern Ethiopia. Our Bayesian occurrence models reveal pronounced stage-specific vulnerabilities: seedling occurrence is highly sensitive to disturbance and browsing pressures, while saplings benefit from larger patch areas yet remain disturbance-prone. Mature individuals exhibit greater resilience, although their long-term stability remains threatened by chronic habitat degradation.

These findings underscore the pressing need for integrated conservation strategies that not only mitigate direct anthropogenic impacts but also enhance the ecological functionality of fragmented patches. Specifically, targeted measures such as controlled grazing regimes, enrichment planting, and microhabitat restoration are essential to bolster early-stage regeneration. Community-based conservation initiatives, aligned with local socio-ecological contexts, will be vital to ensure sustainable stewardship, particularly during critical seasonal transitions from dry to wet periods when seedlings are most vulnerable.

## Data and code availability

The analysis datasets, R scripts (readjusted to suite with our model taken from the book introduction to Win Bugs Bayesian statistics), figures, and tables are publicly available as supplementary material in Get hub

(https://github.com/arayase/church_forest_metacommunity.git)

